# Water-Quality analysis and Fish diversity of Southern West Part of the West-Bengal

**DOI:** 10.1101/2022.06.11.495752

**Authors:** Deep Sankar Chini, Niladri Mondal, Avijit Kar, Ingrid Bunholi, Sourav Singh, Pratik Ghosh, Prasanta Patra, Shampa Patra, Bidhan Chandra Patra

**Affiliations:** Department of Zoology, Vidyasagar University, Midnapore- 721102, West Bengal, India; Department of Biology, Indiana State University, Terre Haute 47809, Indiana, USA; Department of Geography and Environment Management, Vidyasagar University, Midnapore-721102, West Bengal, India

**Keywords:** Water quality, Marine Fish diversity, Seasonal variation, Conservation

## Abstract

Marine fishes are one of the important factors to stabilize the local aquatic ecosystem and regulating socio-economy of local fishermen. In this study we emphasized upon finding out the interrelationship in the water quality, anthropogenic activity, and fish habitat through the 31km stretch of East Medinipur coast in West Bengal, India which is known for its tourist destinations. The study was conducted monthly on different trawler fish landing sites from Dec 2018 to Dec 2021. During this period, we took fish samples and identified them. We took water quality data such as SST, Concentration of Chl-a, Turbidity, and DO for further correlation between the water quality and species diversity. Total 154 numbers of commercially important marine fish species were documented. Among them, 12 species are from Elasmobranch and the rest is Actinopterygii. As per the IUCN database, 13% of the total fish species are under the red list category and 16% of the species showing decreasing population trend. After analyzing the water quality data, we found out that DO, SST, Turbidity, and Chlorophyll-A correlate with the species richness and it also differs during the seasons.

## Introduction

Marine fisheries are an essential supply of food, minerals and other micronutrient resources (Johnson and Welch 2009; Rani 2012). In India, about 18% of fishermen are professionally dependent on marine fisheries sector for their daily livelihood generation (Bhattacharya, Chini et al. 2020). West Bengal including all the southern states and Gujrat have been main areas for marine fish landing stations in India (Sathianandan 2017). Worldwide, marine fish catches remained approximately steady after 1990, while India showed a stable increase up to 3.94 million tons in 2012 (FAO 2018). In the present study we focused on the coastal stretch (about 31 km out of 158 km shoreline) of West Bengal. Recent assessment shows 314 marine fish species are found on the south-western coastal region of West Bengal (Kar, Raut et al. 2017). Continuous unrestrained harvest of marine fishery resources will result in exhaustion of wild population, which leads to considerable impact to aquatic biodiversity and other ecosystem services (Jackson 2008; Christensen 2014). According to FAO report, 33.14% of world ichthyofaunal species are over-exploited and declining very rapidly (FAO 2018). So undoubtedly, the coastal marine ecosystems are becoming vulnerable (already significant changes have been seen in aquatic biota) due to the anthropogenic activities such as changes in water quality, climatic change and overfishing (Jackson, Kirby et al. 2001; McCarthy, Canziani et al. 2001; Metz, Davidson et al. 2001). Abiotic factors play a crucial role for maintaining the species richness and also maintain the natural feeding ground for all marine biota (Bhattacharya, Kar et al. 2018). In focus of the increasing challenges driven by climate change and other natural biotic, abiotic factor sand anthropogenic elements (Parmesan, Burrows et al. 2013; Gilmore, Risi et al. 2018), an enhanced understanding of ichthyofaunal diversity is now in need to support the effective management and development of conservation strategies (Paris, Sherman et al. 2018). Richness and abundance of species will depend upon environmental characteristics such as the SST (Sea Surface Temperature), Chlorophyll (Chl.) Concentration, Dissolve Oxygen (D.O), Turbidity, nutrient load, along with other external factors including invasive species, habitat loss, anthropogenic factors and other pollutants. To find out the present ichthyofaunal diversity, distribution and seasonal variations, we assessed specific parameters of marine water quality along with fish diversity from the different study sites at 31 km stretches of East Medinipur coastline, West Bengal. Study sites are well known for vital marine fishing areas as well as tourism. So, the main objectives of this study were to assess the inter-relationship of water quality, fish diversity, their distribution and role of anthropogenic activity on marine fish diversity in specified area.

## Materials and Methods

### Overview of the study area

The study area is the coastline of West Medinipur, south-western part of West Bengal state, which is enriched by Bay of Bengal. Geographic location in between Latitude 21°36’37.7’’N; Longitude 87°29’10.9’’E to Latitude 21°41’14’’N; Longitude 87°45’19.9’’E comprising about 31 km of coastal shoreline which have been explored during the survey. We have selected the sites for water quality measurement and fish diversity study based on fish harvesting and common landing sites of trawlers from sea. A total of five sampling sites have been selected, which are (1) Old Digha, (2) Digha Mohana, (3) Shankarpur, (4) Tajpurand (5) Mandarmani. Fish sampling and water quality assessment have been done in regular survey-based mode (Coles, Wortley et al. 1985) (**Fig. 1**).

**Fig. 1:**
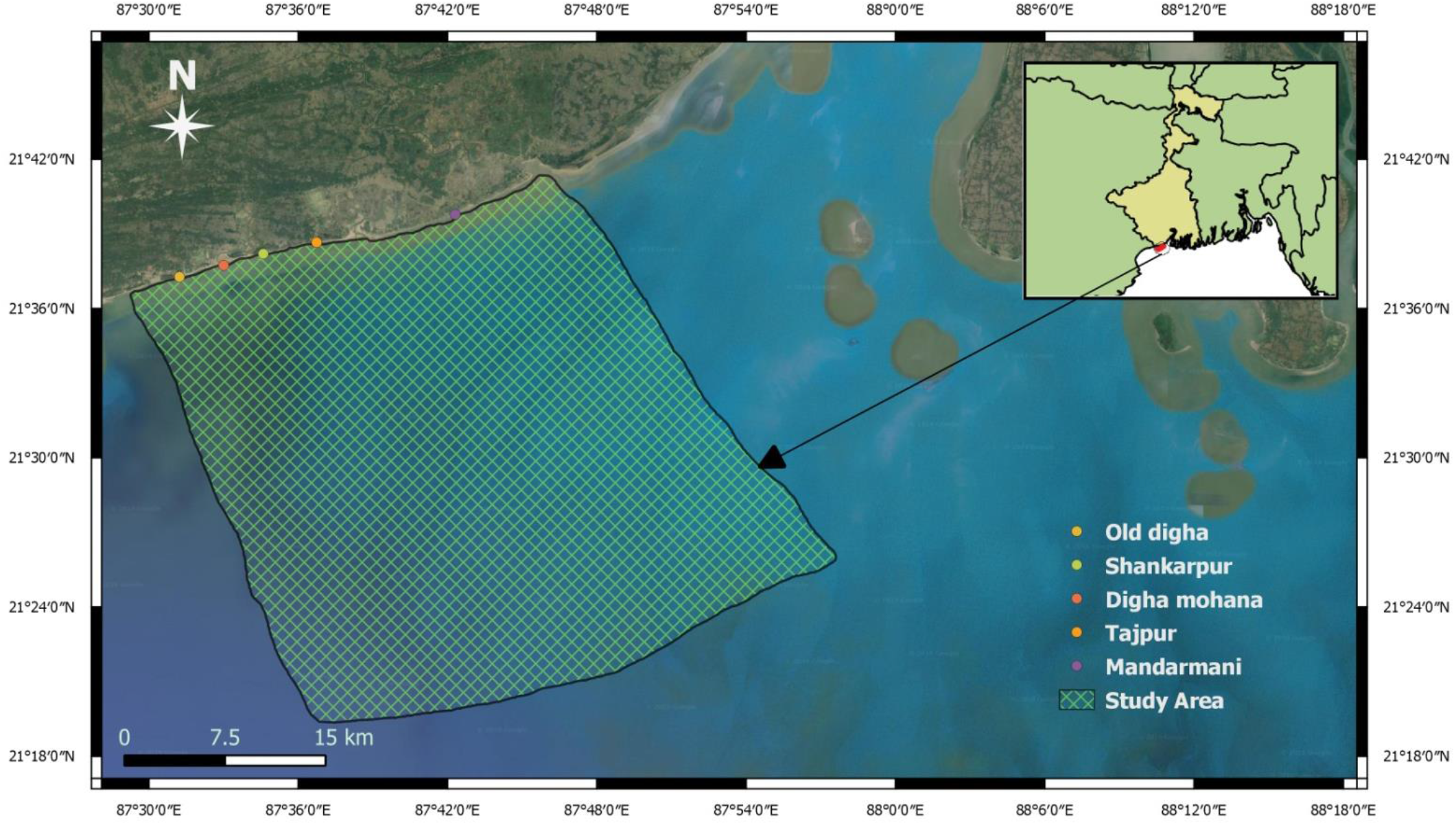
Selected study sites along the West Medinipur coastline.

### In Situ Sampling of Specimens

The specified fishing areas are not only very popular for eco-tourism but also well known for marine fishing through mechanized, non-mechanized fishing boats and trawlers. Each sampling site was surveyed once every month during the study phase of three years (December 2018 to December 2021). During the study period, we accompanied the fishermen on their fishing and early morning inspection of the fish markets for on field data collection. These fishing boats are equipped with gill nets with 9-10 cm mesh size as per West Bengal governmental standard regulation (Edwin 2018). Around 31 km^2^ of marine catchment areas has been covered by this study. After data collection, all fishes have been taxonomically identified following standard references (Talwar and Jhingran 1991). In some circumstances, the several fish specimens (from fish landing stations) were brought into the laboratory in 4% to 6% formaldehyde solution for further taxonomic standardization and preservation. After validation and conservation status assessment of the samples through FishBase ver. 02/2022 (FROESE 2009), WoRMS and IUCN ver. 03/2021 (Mees, Boxshall et al. 2015; IUCN 2021), a checklist of ichthyofauna was prepared. We also did the evaluation of population trend for each species along with their IUCN status and regional occurrence of them in each sampling site.

### Water Parameters analysis

During *In Situ* sampling, five water parameters (Sea surface temperature, concentration of Chlorophyll-a (Chl-a), Turbidity and Dissolved oxygen) were monitored in monthly intervals. Sea surface temperature (SST), DO and Turbidity were calculated by using Systronics Water Analyzer 371 in five sampling area through random sampling in each site (the mean value was taken during the calculation). Concentration of Chl-a was randomly measured through YSI 6025 Chlorophyll Sensor in specified zones (Stumpner, Yin et al. 2022).

### SST and Chlorophyll-a concentration mapping of the study site

We used NOAA’s ERDDAP server data and plotted the SST and Chl-a data from Dec 2018 to Dec 2021 in monthly intervals using R statistical software and map data package (Ihaka and Gentleman 1996; Thomas, Vaughan et al. 2013; Deckmyn 2018). This server generates data of relative SST and Chlorophyll concentration across the globe. We have used the 4km data from the server for our study.

### Statistical analysis

Species richness was calculated using Vegan package on R 4.12 version (Crawley 2012; Oksanen, Blanchet et al. 2013). Correlation between species richness and each water parameter was also calculated using factoextra package on R (Kassambara and Mundt 2017). Species abundance along with number of species were also calculated using iNext package (Hsieh, Ma et al. 2016) on R 4.12 (CoreTeam2021). Every seasonal variety of species richness was also calculated to determine the variability during seasonal changes. IUCN red listed species were plotted against a heat map to see their seasonal availability of by catch during the study period.

## Results and Discussions

### Fish checklist and diversity

A total of 154 marine fish species were identified by this survey. Among them, 10% species belong to the Elasmobranch group while the other 90% comprise of commercially important ray-finned species (supp1). Most of the recorded fishes have a high economic value for being used as food, byproducts as well as ornaments in the region. The fishes belong to the order Perciformes followed by Clupeiiformes and Siluriiformes) and 58 families (notable families are Clupeiidae, Engraulidae, Sciaenidae, Mugilidae and Ariidae). During the commercial fish catches fish landing showed maximum catchment was done during the month of July to September and lowest in March to April (In West Bengal 15^th^ April to 14^th^ June is ban period for fishing due to their spawning). Around 13% of the identified species are listed in the IUCN Red List (IUCN, 2019) (**Fig. 2a**). The population trends are stable (8%), decreasing (16%) and rest of the recorded species are under unknown (75%) status **(Fig. 2.b)**.

**Fig 2.a:**
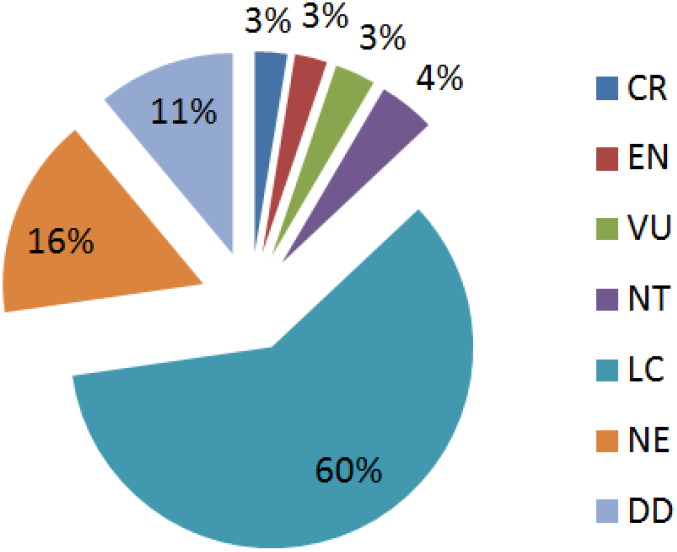
IUCN (ver. 2021-3) Categorization.

**Fig 2.b:**
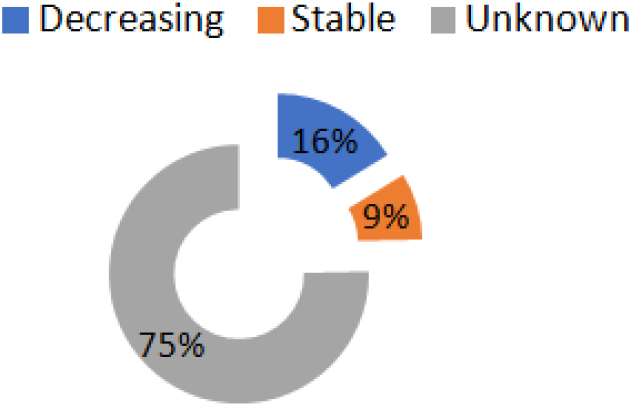
Population trend based on IUCN Ver. 2021-3.

The list of 20 species which are under CR, EN, VU and NT status points out with their present population trends **(Table 1)**. Depending on the seasonal variation of availability of concerned 20 IUCN red listed fish species are represented by the matrix plot **(Fig. 3)**. The plot also explains that *Cacrhahinus limbatus, Carchahinus sorrah, Rhizoprionodon djiddensis* were rarely seen during the study period whereas *Glaucotegus granulatus, Eleutheronema tetradactylum, Omnok bimaculatus* were found frequently. Majority of the fishes were reported during the season of monsoon and post-monsoon period as those time the highest number of fish were caught for food (**Fig. 3**). which could be alarming and specific conservation action plan is needed for them. Species *Anguilla bengalensi* and *Scombeomorus commerson* and *Tenualosa toli* have been the most affected ones as their occurrence number has been less in the last three sampling period.

**Table.1:**
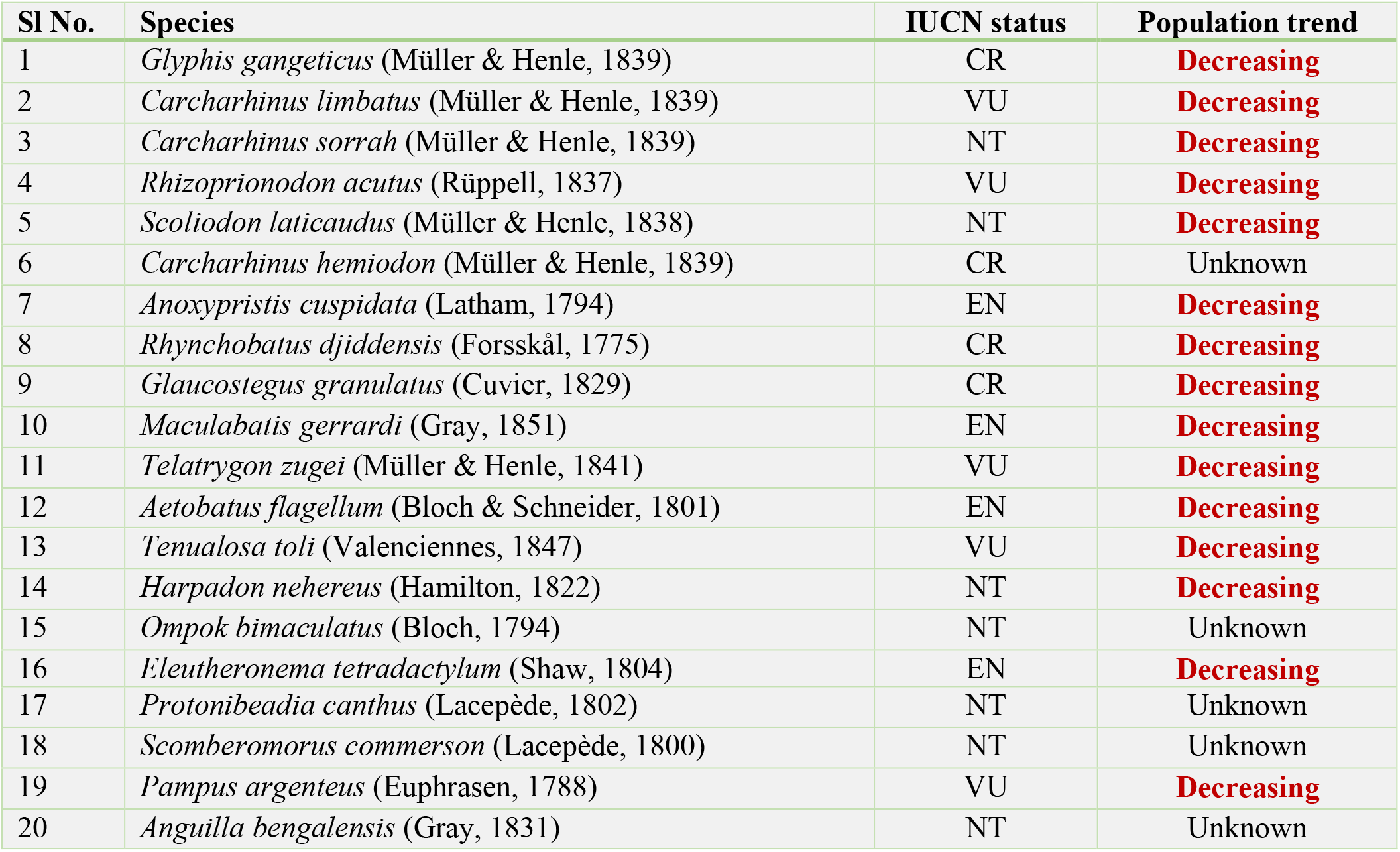
List of IUCN red listed elasmobranch and ray-finned species along with their population trend.

**Fig. 3:**
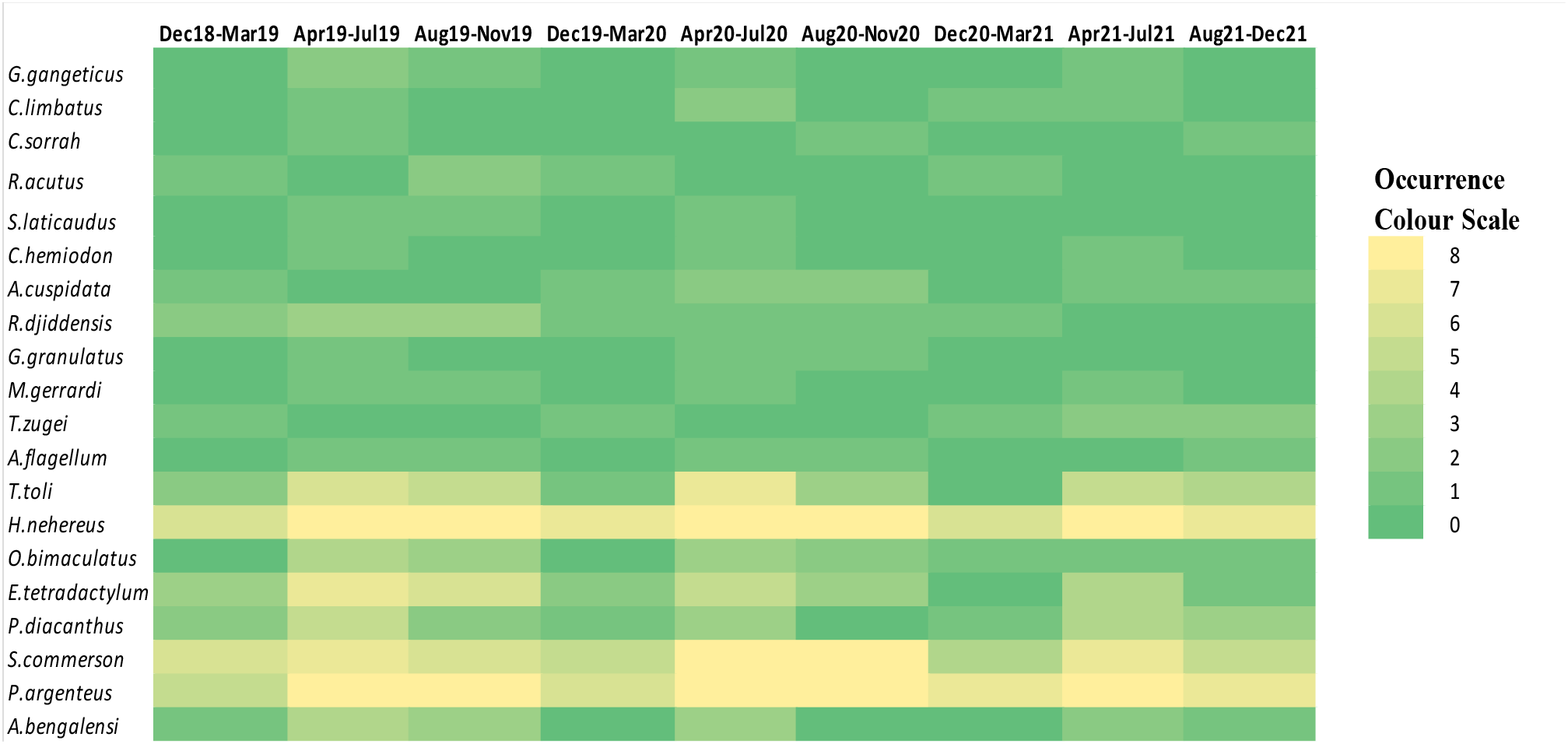
The occurrence matrix plot of IUCN relisted Elasmobranch and Actinoptery species from study sites based on seasonal variation.

Total 6,678 number of fish samples was considered from all the predetermined sampling sites for diversity indices data analysis. The maximum numbers of fishes were collected from Shankarpur (29.9%), followed by Digha Mohana (28.4%), Mandarmani (18.6%), Tajpur (14.4%) and Old Digha (8.7%) sampling sites. It also confirms that getting more tourists and getting high anthropogenic load, Old Digha site shows less species diversity (63 species) whereas the highest species diversity was found at Digha-Mohana (107 species) and Shankarpur study sites (102 species).The number of species and sample size was plotted for each of these selected areas, which shows high amount of fish catches and availability in those areas (**Fig. 4a**). We selected 4 seasons to show the species richness according to those seasons and fish availability data (**Fig. 4b)**. Generally, fish catchment was low during the winter season. Therefore, we found a smaller number of species in fish markets during the winter season and the beginning of post-monsoon due to having a State-wide ban on fishing for fish spawning period.

**Fig. 4:**
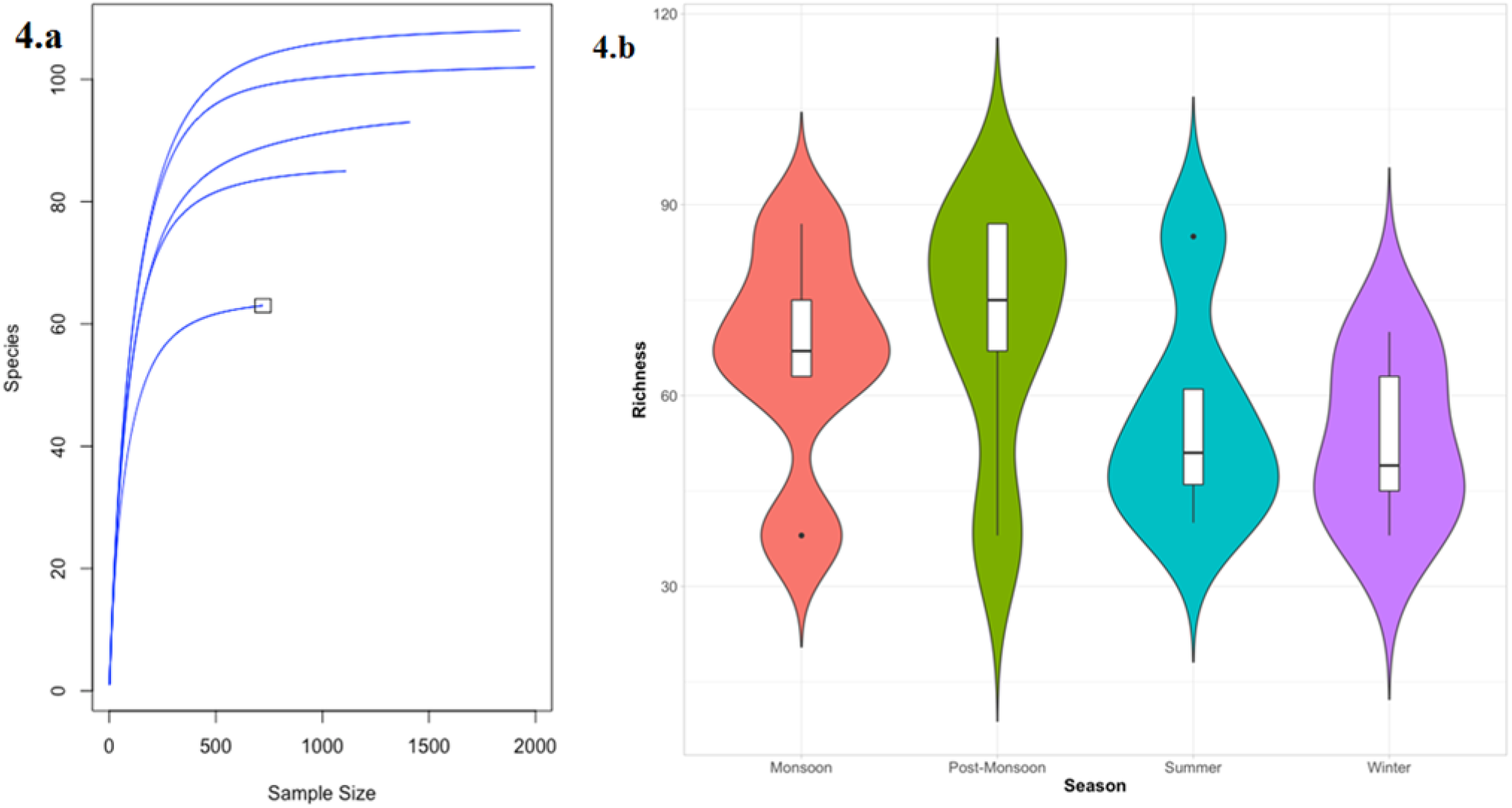
Fish sample estimation and seasonal variety of species richness plot.

### Water quality assessment and GIS mapping

We have estimated the monthly SST and VIIRS monthly Chl-a data of our study site. Estimated SST data shows relatively lower temperature on the shore region which have allowed fish gathering and species abundance throughout the year (**Fig. 5a**). Monthly chlorophyll-a concentration data shows the relative high amount of chlorophyll concentration close to the shoreline (**Fig. 5b**), which might be correlated with the high accumulation of microbiome and phytoplankton density, suitable for fish nutrition.

**Fig.5.**
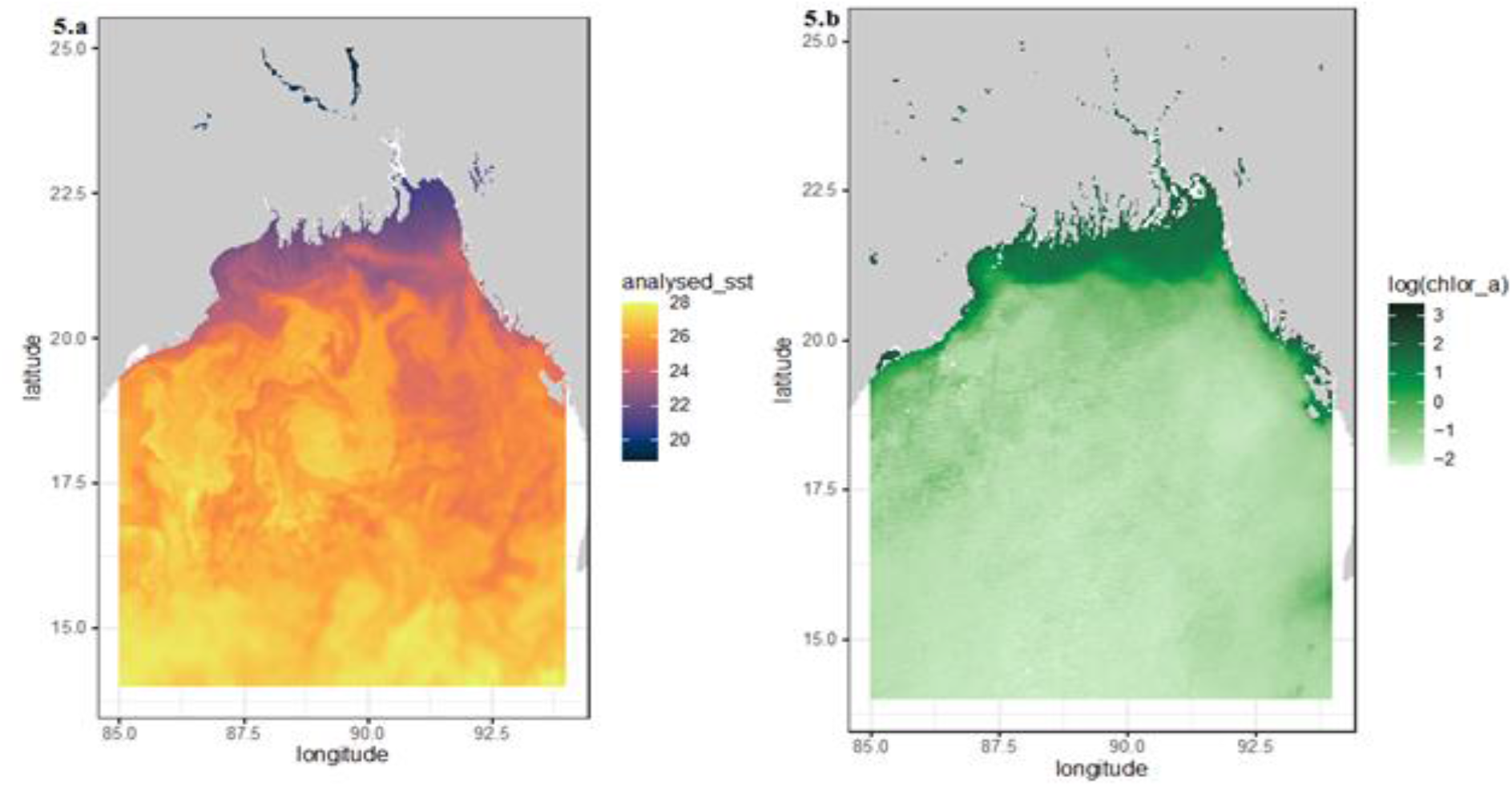
Analyzed Seas Surface Temperature (SST) and Chlorophyll-a concentration map of the Bay of Bengal.

DO variation map depending on the season shows high DO during winter season and Monsoon season on the Digha-Mohana site(**fig:6**). While lower DO has been detected on the Shankarpur and Old Digha sites.

**Fig:6.**
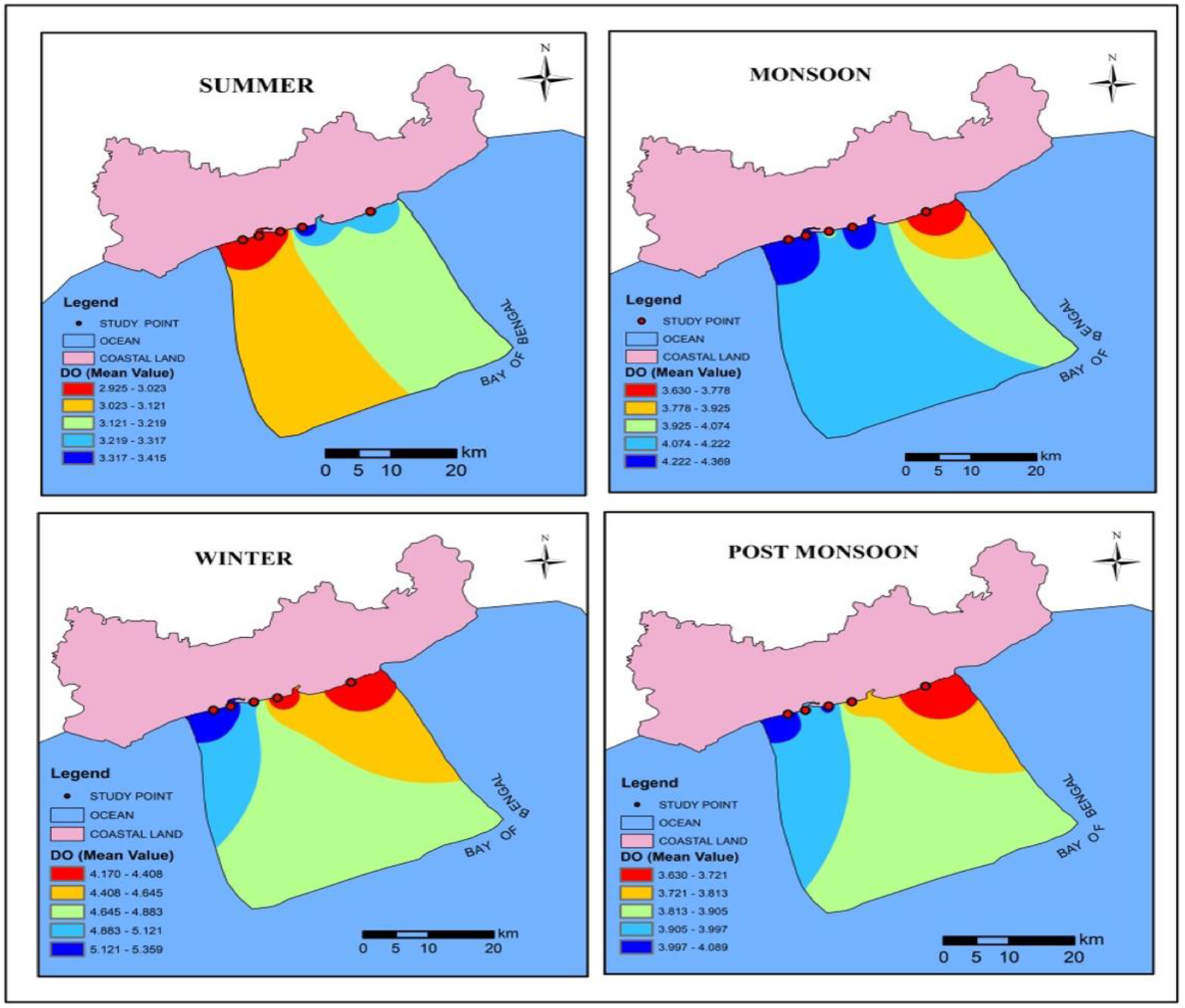
Analyzed Dissolved Oxygen based on Seasonal Changes of the study site.

Turbidity of the study sites has been changed due to seasonal variation, highest turbidity has been analyzed on the Tazpur and Mandarmani area(fig:7).

**Fig:7.**
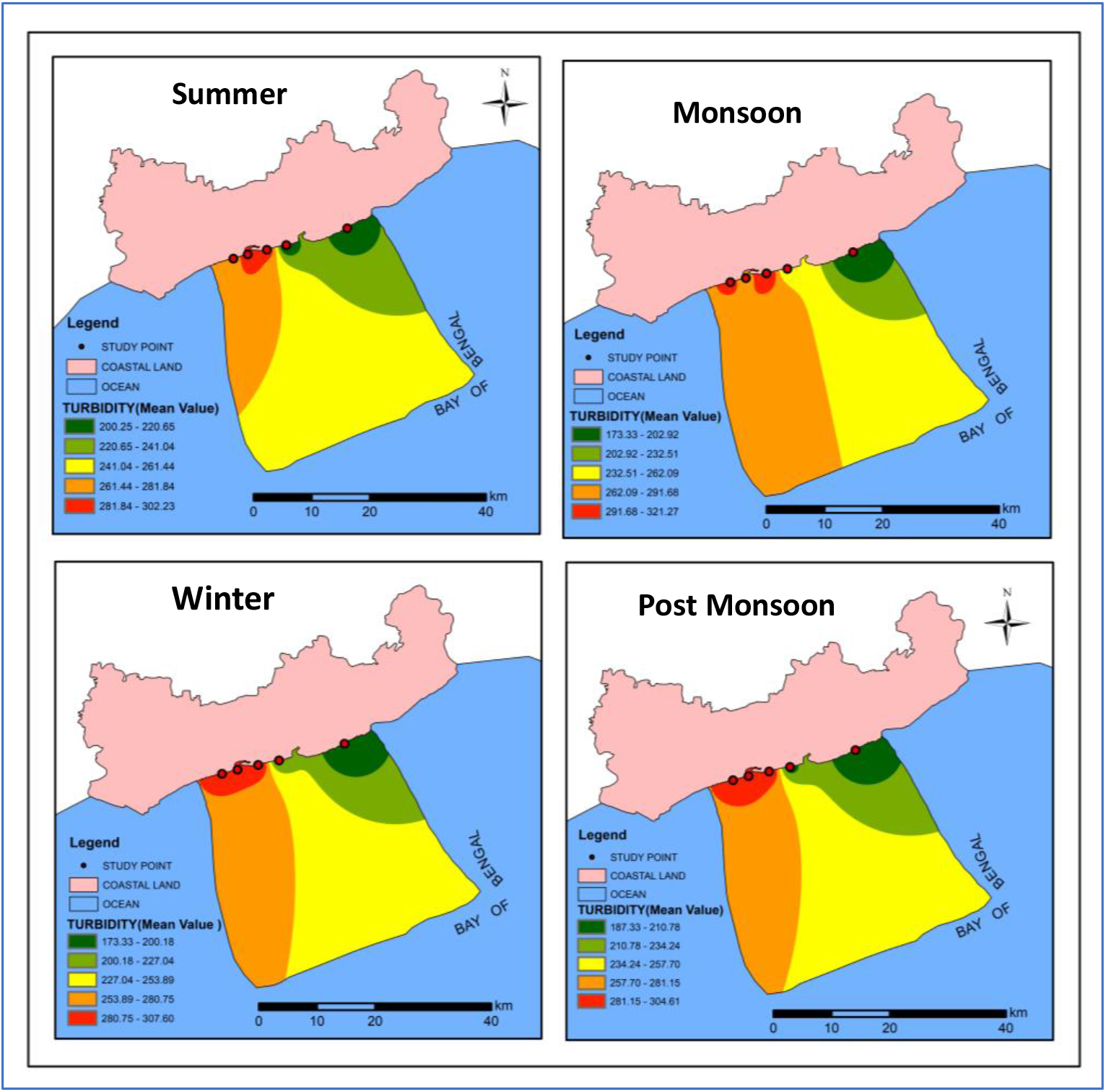
Distribution of turbidity based on seasonal variation on the study sites.

### Water quality analysis of selected sites in seasonal changes

Sea Surface Temperature fluctuates throughout the year as in summer the SST reaches to highest while in monsoon and post monsoon period the temperature is moderate which reflected on the species richness (**Fig. 8a**). Species richness affects during increasing anthropogenic activity is seen on the coastal areas (McKinney 2008). As we can see the tourist regions including Digha and Mandarmani has significant decrease of species richness (**Fig. 8b**).

**Fig.8.**
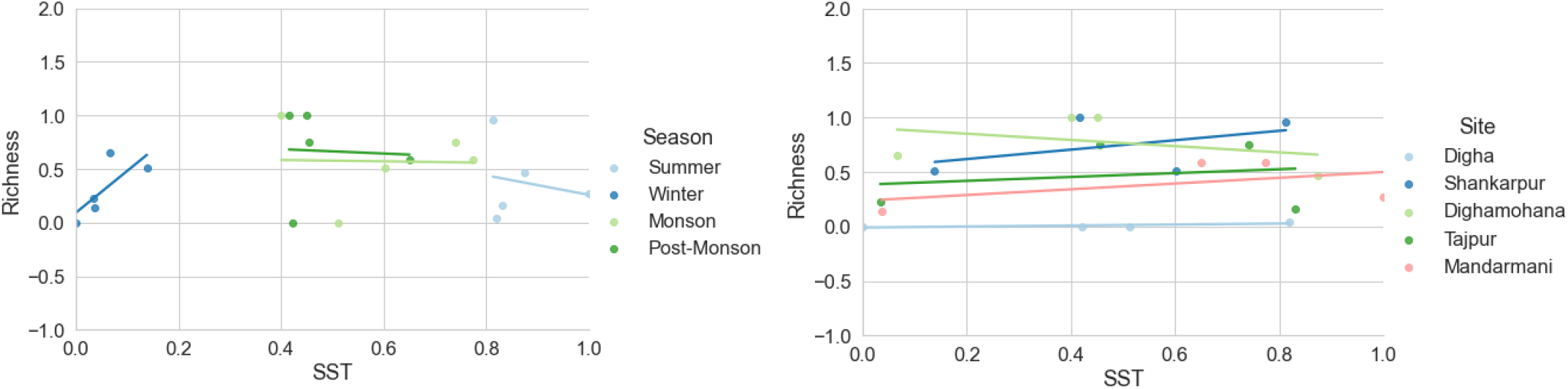
Correlation of SST along with seasonal variety and sampling sites.

Chlorophyll concentration of the coastal area changes by season (Boyce, Dowd et al. 2014). During the monsoon and post monsoon period, large outflow of river water into the estuaries and then final intake into the ocean releases huge microbiome and organic matter resulting in increasing chlorophyll concentration (**Fig. 9a**). Chlorophyll concentration stayed low during the less productive winter season. Higher chlorophyll concentration results in less species richness while the study sites also showed different range. Site Shankarpur and Digha-mohana showed the highest Chlorophyll concentration as well the highest fish species richness due to containing both the estuaries and the most trawler landing sites among the selected regions (**Fig. 9b**).

**Fig. 9.**
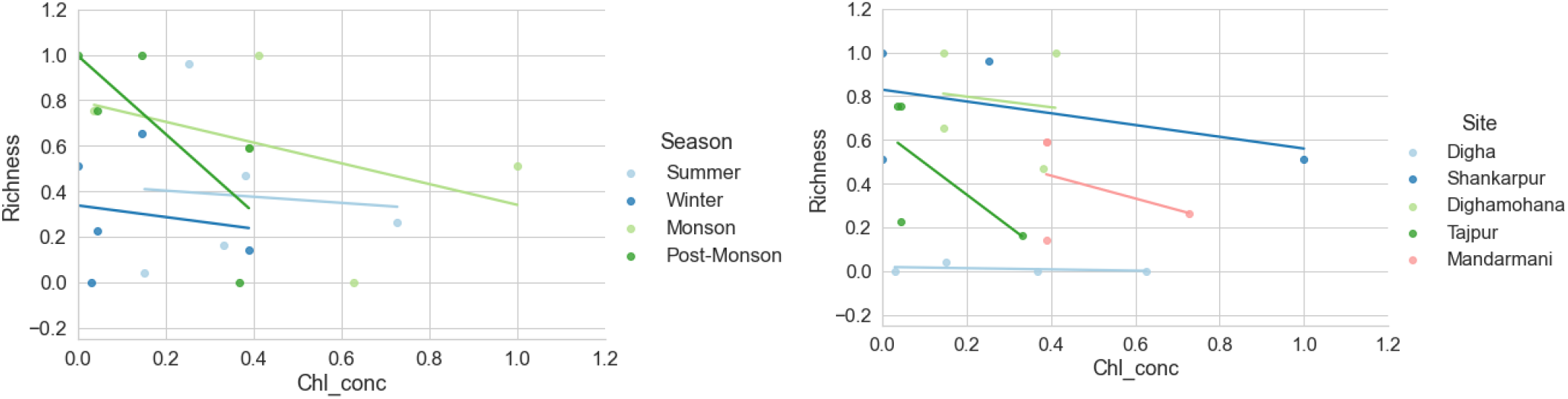
Correlation of Chlorophyll -a concentration in respect to seasonal variety and Sampling sites.

Reduced oxygen due to less nutrient loading in water can be dangerous for fish population (Joos, Plattner et al. 2003). Most of the algal bloom of the oxygen producing water plant growth is hindered when there is cloudy condition or when there is much more anthropogenic activity which usually in the summer or in the winter (**Fig. 10a**). Species richness positively correlates with the DO that is why species richness is higher during monsoon and post monsoon season among all the season. Species richness also fluctuates on the study sites depending on the DO, site Digha is the most affected one in case of species most of the time associated with tourist seasons (Roy ; Pitchaikani, Kadharsha et al. 2016; Roy and Shamim 2020). While being the major fish landing centers; Digha-mohana and Shankarpur show more species richness (**Fig. 10b**).

**Fig. 10.**
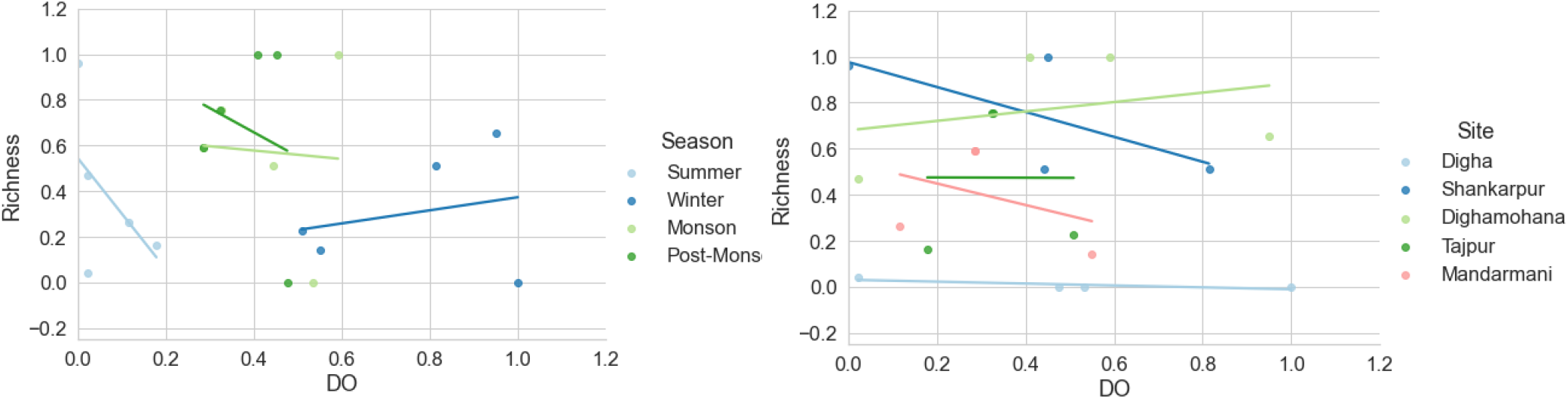
Correlation of Dissolved Oxygen in respect to seasonal variety and Sampling sites.

Turbidity of the sea water depends on various factors including eutrophication, sedimentary runoff, dissolved colored organic matter along with high number of phytoplankton (Pitchaikani, Kadharsha et al. 2016). Even though turbidity sometimes may affect the fish diversity and species richness (mostly in the sea water where greater portion of coral reefs are present), our study sites have shown positive relationship with turbidity. A huge intake of river water from the estuaries and a seasonal max turbidity can be seen during monsoon and post monsoon period on the study sites where they also attain high species richness (**Fig. 11a**). Turbidity can also be elevated by anthropogenic activity by resurfacing the water during the tourist seasons, that is why the turbidity is on the high rise at the Digha site (**Fig. 11b**) even though the species richness is not that much higher on the region. Fish-landing sites Shankarpur and Digha-mohana both having high species richness and elevated turbidity due to those factors (**Fig. 11b**).

**Fig.11.**
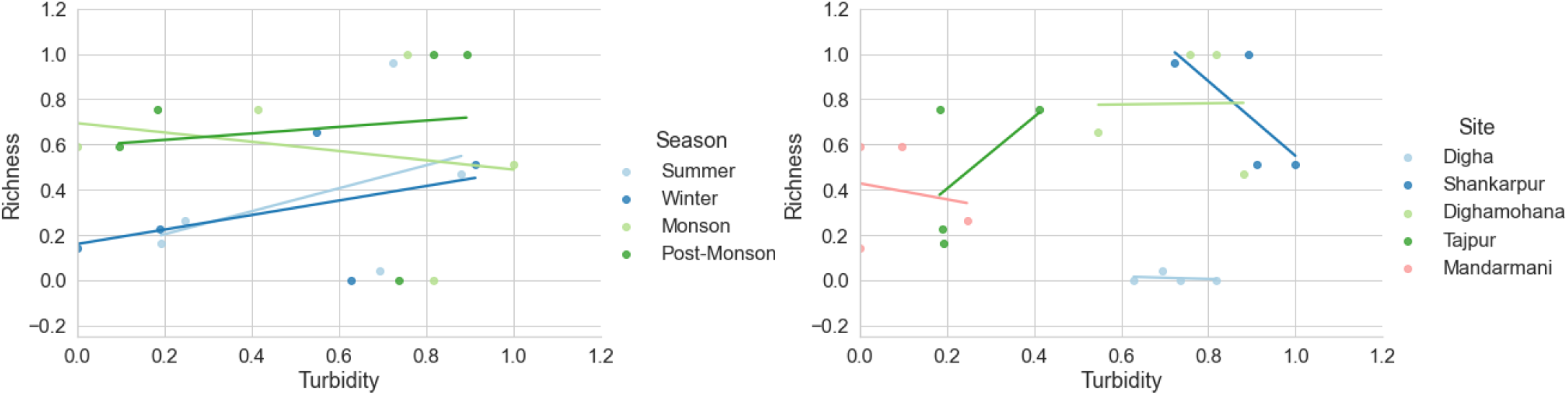
Correlation of Turbidity c in respect to seasonal variety and Sampling sites.

## Conclusion

Present study shows the changes in water quality and the species richness depending on the season. Study sites where the anthropogenic activity is higher than the other sites show relatively less richness. We found 20 species which are IUCN red listed and 15 out of those 20 species populations are declining. This study presents the current status of the fish which are economically important for the coastal areas of west Bengal. This will allow other researchers to start working and putting more emphasis on the southern coastal region of West Bengal which have not been highlighted in the past few years. Most elasmobranchs we found during the survey show population decline. Those species are also important to stabilize the aquatic environments, to gather more knowledge about these red listed species and their habitat status, proper analysis of habitat and by catch information will be essential. Our next goal is to get by catch data of those species which are not commercially valuable but get caught and cause the decline of their population.

Considering the socioeconomic significance of the Bay of Bengal coastal regions for tourism and fishing for livelihood and commercial purposes, water quality analysis and fish assemblage study is important for conservation. This study highlights the importance of assessing pollution within coastal regions in relation to the species richness and changes in water quality on the seasonal basis. The future studies should help to ensure the marine fish community stabilization, stock assessment and socioeconomic upliftment of fisher folks. The work also finds out the IUCN relisted Elasmobranch and Actinoptery which are harvested in uncontrolled manner. So, fish assemblages could be used as ecosystem functioning indicators, and certain fish populations should be permanent monitored before their extinction.

## Author contribution

D.S.C and N.M. and B.P. wrote the manuscript with input from all the authors. A.K, P.G and P.P helped during the data collection. I.B., S.P. and S.S. analyzed the data and visualize it.

## Acknowledgements

Authors are thankful to WBHESTB, Govt. of West Bengal (File no.: ST/S & T/1G-13/2017) for sanctioning the research project as well as Science and Engineering Research Board, Department of Science and Technology, Government of India for financial assistance to carry out the research work (Project file No Ref. No.PDF/2016/001776). Authors also like to thank the Council of Scientific and Industrial Research (CSIR) sponsored Junior Research Fellowship (File No.-09/599(0087)/2019-EMR-1) to P.G.

