## Supplemental Table 1 for "Water-Quality analysis and Fish diversity of Southern West Part of the West-Bengal"

*Check list of Marine fish diversity in the study area with taxonomic position, IUCN status and population trend*

| SL. No. | Sub-Class | Order | Family | Name | IUCN Ver. 2021-3 | Population trends |
|---|---|---|---|---|---|---|
| 1 | Elasmobranchii | Carcharhiniformes | Carcharhinidae | *Glyphis gangeticus* (Müller & Henle, 1839) | CR | **Decreasing** |
| 2 | | | | *Carcharhinus limbatus* (Müller & Henle, 1839) | VU | **Decreasing** |
| 3 | | | | *Carcharhinus sorrah* (Müller & Henle, 1839) | NT | **Decreasing** |
| 4 | | | | *Rhizoprionodon acutus* (Rüppell, 1837) | VU | **Decreasing** |
| 5 | | | | *Scoliodon laticaudus* (Müller & Henle, 1838) | NT | **Decreasing** |
| 6 | | | | *Carcharhinus hemiodon* (Müller & Henle, 1839) | CR | Unknown |
| 7 | | Rhinopristiformes | Pristidae | *Anoxypristis cuspidata* (Latham, 1794) | EN | **Decreasing** |
| 8 | | | Rhinidae | *Rhynchobatus djiddensis* (Forsskål, 1775) | CR | **Decreasing** |
| 9 | | | Glaucostegidae | *Glaucostegus granulatus* (Cuvier, 1829) | CR | **Decreasing** |
| 10 | | Myliobatiformes | Dasyatidae | *Maculabatis gerrardi* (Gray, 1851) | EN | **Decreasing** |
| 11 | | | | *Telatrygon zugei* (Müller & Henle, 1841) | VU | **Decreasing** |
| 12 | | | Myliobatidae | *Aetobatus flagellum* (Bloch & Schneider, 1801) | EN | **Decreasing** |
| 13 | Actinopterygii | Clupeiformes | Clupeidae | *Tenualosa ilisha* (Hamilton, 1822) | LC | **Decreasing** |
| 14 | | | | *Tenualosa toli* (Valenciennes, 1847) | VU | **Decreasing** |
| 15 | | | | *Hilsa kelee* (Cuvier, 1829) | LC | **Stable** |
| 16 | | | | *Sardinella brachysoma* (Bleeker, 1852) | LC | Unknown |
| 17 | | | | *Sardinella fimbriata* (Valenciennes, 1847) | LC | Unknown |
| 18 | | | | *Sardinella gibbosa* (Bleeker, 1849) | LC | Unknown |
| 19 | | | | *Gonialosa manmina* (Hamilton, 1822) | LC | Unknown |
| 20 | | | | *Anodontostoma chacunda* (Hamilton, 1822) | LC | Unknown |
| 21 | | | | *Nematalosa nasus* (Bloch, 1795) | LC | Unknown |
| 22 | | | | *Escualosa thoracata* (Valenciennes, 1847) | LC | Unknown |
| 23 | | | | *Amblygaster leiogaster* (Valenciennes, 1847) | LC | Unknown |
| 24 | | | | *Sardinella albella* (Valenciennes, 1847) | LC | Unknown |
| 25 | | | Pristigasteridae | *Ilisha kampeni* (Weber & de Beaufort, 1913) | LC | Unknown |
| 26 | | | | *Ilisha melastoma* (Bloch & Schneider, 1801) | LC | Unknown |
| 27 | | | | *Raconda russeliana* (Gray, 1831) | LC | Unknown |
| 28 | | | | *Pellona ditchela* (Valenciennes, 1847) | LC | Unknown |
| 29 | | | Engraulidae | *Coilia dussumieri* (Valenciennes, 1848) | LC | **Stable** |
| 30 | | | | *Coilia ramcarati* (Hamilton, 1822) | DD | Unknown |
| 31 | | | | *Coilia reynaldi* (Valenciennes, 1848) | LC | **Stable** |
| 32 | | | | *Setipinna phasa* (Hamilton, 1822) | LC | Unknown |
| 33 | | | | *Setipinna tenuifilis* (Valenciennes, 1848) | DD | Unknown |
| 34 | | | | *Setipinna taty* (Valenciennes, 1848) | LC | Unknown |
| 35 | | | | *Stolephorus indicus* (van Hasselt, 1823) | LC | Unknown |
| 36 | | | | *Stolephorus commersonnii* (Lacepède, 1803) | LC | Unknown |
| 37 | | | | *Thryssa malabarica* (Bloch, 1795) | DD | Unknown |
| 38 | | | | *Thryssa purava* (Hamilton, 1822) | DD | Unknown |
| 39 | | | | *Thryssa hamiltonii* (Gray, 1835) | LC | Unknown |

*Check list of Marine fish diversity in the study area with taxonomic position, IUCN status and population trend*

| | | | | | |
|---|---|---|---|---|---|
| 40 | | Chirocentridae | *Chirocentrus dorab* (Forsskål, 1775) | LC | Unknown |
| 41 | | Dussumieriidae | *Dussumieria acuta* (Valenciennes, 1847) | LC | Unknown |
| 42 | Elopiformes | Megalopidae | *Megalops cyprinoides* (Broussonet, 1782) | DD | Unknown |
| 43 | Aulopiformes | Synodontidae | *Harpadon nehereus* (Hamilton, 1822) | NT | **Decreasing** |
| 44 | Siluriformes | Ariidae | *Arius jella* (Day, 1877) | NE | Unknown |
| 45 | | | *Hexanematichthys sagor* (Hamilton, 1822) | NE | Unknown |
| 46 | | | *Plicofollis layardi* (Günther, 1866) | NE | Unknown |
| 47 | | | *Nemapteryx caelata* (Valenciennes, 1840) | NE | Unknown |
| 48 | | | *Osteogeneiosus militaris* (Linnaeus, 1758) | NE | Unknown |
| 49 | | | *Plicofollis platystomus* (Day, 1877) | LC | Unknown |
| 50 | | | *Plicofollis dussumieri* (Valenciennes, 1840) | LC | **Stable** |
| 51 | | | *Sciades sona* (Hamilton, 1822) | NE | Unknown |
| 52 | | Plotosidae | *Plotosus canius* (Hamilton, 1822) | NE | Unknown |
| 53 | | Silurida | *Ompok bimaculatus* (Bloch, 1794) | NT | Unknown |
| 54 | | Schilbeidae | *Eutropiichthys vacha* (Hamilton, 1822) | LC | **Decreasing** |
| 55 | | | *Clupisoma garua* (Hamilton, 1822) | LC | **Decreasing** |
| 56 | | | *Silonia silondia* (Hamilton, 1822) | LC | Unknown |
| 57 | | Pangasiidae | *Pangasius pangasius* (Hamilton, 1822) | LC | **Decreasing** |
| 58 | Anguilliformes | Anguillidae | *Anguilla bengalensis* (Gray, 1831) | NT | Unknown |
| 59 | | Moringuidae | *Moringua raitaborua* (Hamilton, 1822) | NE | Unknown |
| 60 | | Muraenidae | *Gymnothorax pictus* (Ahl, 1789) | LC | Unknown |
| 61 | | Muraenesocidae | *Congresox talabon* (Cuvier, 1829) | NE | Unknown |
| 62 | | | *Congresox talabonoides* (Bleeker, 1853) | NE | Unknown |
| 63 | | Ophichthidae | *Pisodonophis boro* (Hamilton, 1822) | LC | Unknown |
| 64 | | | *Lamnostoma orientalis* (McClelland, 1844) | LC | Unknown |
| 65 | Beloniformes | Belonidae | *Strongylura strongylura* (van Hasselt, 1823) | NE | Unknown |
| 66 | | Hemiramphidae | *Hyporhamphus quoyi* (Valenciennes, 1847) | NE | Unknown |
| 67 | | | *Hyporhamphus affinis* (Günther, 1866) | NE | Unknown |
| 68 | | | *Rhynchorhamphus georgii* (Valenciennes, 1847) | LC | Unknown |
| 69 | | Zenarchopteridae | *Zenarchopterus ectuntio* (Hamilton, 1822) | LC | Unknown |
| 70 | Mugiliformes | Mugilidae | *Planiliza parsia* (Hamilton, 1822) | NE | Unknown |
| 71 | | | *Planiliza planiceps* (Valenciennes, 1836) | LC | Unknown |
| 72 | | | *Planiliza macrolepis* (Smith, 1846) | LC | **Stable** |
| 73 | | | *Osteomugil cunnesius* (Valenciennes, 1836) | NE | Unknown |
| 74 | | | *Planiliza subviridis* (Valenciennes, 1836) | LC | Unknown |
| 75 | | | *Mugil cephalus* (Linnaeus, 1758) | LC | **Stable** |
| 76 | | | *Ellochelon vaigiensis* (Quoy & Gaimard, 1825) | LC | Unknown |
| 77 | | | *Rhinomugil corsula* (Hamilton, 1822) | LC | Unknown |
| 78 | Perciformes | Polynemidae | *Eleutheronema tetradactylum* (Shaw, 1804) | EN | **Decreasing** |
| 79 | | | *Polydactylus plebeius* (Broussonet, 1782) | NE | Unknown |
| 80 | | | *Polydactylus sextarius* (Bloch & Schneider, 1801) | NE | Unknown |

*Check list of Marine fish diversity in the study area with taxonomic position, IUCN status and population trend*

| | | | | | |
|---|---|---|---|---|---|
| 81 | | | *Polynemus paradiseus* (Linnaeus, 1758) | LC | **Decreasing** |
| 82 | | | *Leptomelanosoma indicum* (Shaw, 1804) | NE | Unknown |
| 83 | | Latidae | *Lates calcarifer* (Bloch, 1790) | LC | Unknown |
| 84 | | Ambassidae | *Ambassis nalua* (Hamilton, 1822) | LC | Unknown |
| 85 | | Terapontidae | *Terapon jarbua* (Forsskål, 1775) | LC | Unknown |
| 86 | | | *Terapon theraps* (Cuvier, 1829) | LC | Unknown |
| 87 | | Sillaginidae | *Sillago sihama* (Forsskål, 1775) | LC | **Stable** |
| 88 | | | *Sillaginopsis panijus* (Hamilton, 1822) | NE | Unknown |
| 89 | | Carangidae | *Megalaspis cordyla* (Linnaeus, 1758) | LC | Unknown |
| 90 | | | *Atropus atropos* (Bloch & Schneider, 1801) | LC | Unknown |
| 91 | | | *Alectis indica* (Rüppell, 1830) | LC | Unknown |
| 92 | | | *Caranx sexfasciatus* (Quoy & Gaimard, 1825) | LC | **Decreasing** |
| 93 | | | *Carangoides malabaricus* (Bloch & Schneider, 1801) | LC | Unknown |
| 94 | | | *Selaroides leptolepis* (Cuvier, 1833) | LC | Unknown |
| 95 | | | *Parastromateus niger* (Bloch, 1795) | LC | **Decreasing** |
| 96 | | Sphyraenidae | *Sphyraena obtusata* (Cuvier, 1829) | LC | **Stable** |
| 97 | | Lactariidae | *Lactarius lactarius* (Bloch & Schneider, 1801) | DD | Unknown |
| 98 | | Menidae | *Mene maculata* (Bloch & Schneider, 1801) | NE | Unknown |
| 99 | | Lutjanidae | *Lutjanus johnii* (Bloch, 1792) | LC | Unknown |
| 100 | | | *Lutjanus indicus* (Allen, White & Erdmann, 2013) | LC | Unknown |
| 101 | | | *Lutjanus malabaricus* (Bloch & Schneider, 1801) | LC | Unknown |
| 102 | | | *Lutjanus russellii* (Bleeker, 1849) | LC | Unknown |
| 103 | | | Lutjanus argentimaculatus (Forsskål, 1775) | LC | Unknown |
| 104 | | Leiognathidae | *Leiognathus equulus* (Forsskål, 1775) | LC | Unknown |
| 105 | | | *Eubleekeria splendens* (Cuvier, 1829) | LC | **Stable** |
| 106 | | Gerreidae | *Gerres filamentosus* (Cuvier, 1829) | LC | Unknown |
| 107 | | | *Gerres setifer* (Hamilton, 1822) | NE | Unknown |
| 108 | | | *Pentaprion longimanus* (Cantor, 1849) | LC | Unknown |
| 109 | | Haemulidae | *Pomadasys maculatus* (Bloch, 1793) | LC | Unknown |
| 110 | | Sciaenidae | *Macrospinosa cuja* (Hamilton, 1822) | DD | Unknown |
| 111 | | | *Dendrophysa russelii* (Cuvier, 1829) | LC | Unknown |
| 112 | | | *Otolithoides biauritus* (Cantor, 1849) | DD | **Decreasing** |
| 113 | | | *Nibea soldado* (Lacepède, 1802) | LC | Unknown |
| 114 | | | *Protonibea diacanthus* (Lacepède, 1802) | **NT** | **Decreasing** |
| 115 | | | *Pterotolithus maculatus* (Cuvier, 1830) | LC | Unknown |
| 116 | | | *Johnius belangerii* (Cuvier, 1830) | LC | Unknown |
| 117 | | | *Johnius coitor* (Hamilton, 1822) | LC | Unknown |
| 118 | | | *Johnius borneensis* (Bleeker, 1851) | LC | Unknown |
| 119 | | | *Otolithoides pama* (Hamilton, 1822) | DD | Unknown |
| 120 | | Drepaneidae | *Drepane punctata* (Linnaeus, 1758) | LC | **Stable** |

*Check list of Marine fish diversity in the study area with taxonomic position, IUCN status and population trend*

| | | | | | |
|---|---|---|---|---|---|
| 121 | | Trichiuridae | *Eupleurogrammus muticus* (Gray, 1831) | DD | Unknown |
| 122 | | | *Lepturacanthus savala* (Cuvier, 1829) | NE | Unknown |
| 123 | | | *Trichiurus lepturus* (Linnaeus, 1758) | **LC** | **Stable** |
| 124 | | | *Lepturacanthus pantului* (Gupta, 1966) | DD | Unknown |
| 125 | | | *Eupleurogrammus muticus* (Gray, 1831) | DD | Unknown |
| 126 | | Scombridae | *Euthynnus affinis* (Cantor, 1849) | **LC** | Unknown |
| 127 | | | *Scomberomorus commerson* (Lacepède, 1800) | **NT** | **Decreasing** |
| 128 | | | *Rastrelliger kanagurta* (Cuvier, 1816) | DD | Unknown |
| 129 | | | *Auxis thazard* (Lacepède, 1800) | **LC** | Stable |
| 130 | | | *Scomberomorus guttatus* (Bloch & Schneider, 1801) | DD | Unknown |
| 131 | | Stromateidae | *Pampus argenteus* (Euphrasen, 1788) | **VU** | **Decreasing** |
| 132 | | | *Pampus chinensis* (Euphrasen, 1788) | NE | Unknown |
| 133 | | Gobiidae | *Pseudapocryptes elongatus* (Cuvier, 1816) | **LC** | Unknown |
| 134 | | | *Apocryptes bato* (Hamilton, 1822) | **LC** | Unknown |
| 135 | | | *Scartelaos histophorus* (Valenciennes, 1837) | **LC** | Unknown |
| 136 | | | *Boleophthalmus boddarti* (Pallas, 1770) | **LC** | Unknown |
| 137 | | | *Trypauchen vagina* (Bloch & Schneider, 1801) | **LC** | Unknown |
| 138 | | | *Odontamblyopus rubicundus* (Hamilton, 1822) | **LC** | Unknown |
| 139 | | Sparidae | *Acanthopagrus berda* (Forsskål, 1775) | **LC** | Unknown |
| 140 | | | *Acanthopagrus longispinnis* (Valenciennes, 1830) | DD | Unknown |
| 141 | | Nemipteridae | *Nemipterus peronii* (Valenciennes, 1830) | **LC** | Unknown |
| 142 | | Mullidae | *Upeneus sulphureus* (Cuvier, 1829) | **LC** | **Stable** |
| 143 | | Scatophagidae | *Scatophagus argus* (Linnaeus, 1766) | **LC** | Unknown |
| 144 | | Istiophoridae | *Istiophorus platypterus* (Shaw, 1792) | **LC** | Unknown |
| 145 | Pleuronectiformes | Psettodidae | *Psettodes erumei* (Bloch & Schneider, 1801) | DD | Unknown |
| 146 | | Paralichthyidae | *Pseudorhombus arsius* (Hamilton, 1822) | **LC** | Unknown |
| 147 | | Cynoglossidae | *Paraplagusia bilineata* (Bloch, 1787) | **LC** | Unknown |
| 148 | | | *Cynoglossus cynoglossus* (Hamilton, 1822) | **LC** | Unknown |
| 149 | | Soleidae | *Synaptura albomaculata* (Kaup, 1858) | **LC** | Unknown |
| 150 | | | *Brachirus pan* (Hamilton, 1822) | **LC** | Unknown |
| 151 | | | *Synaptura commersonnii* (Lacepède, 1802) | **LC** | Unknown |
| 152 | Scorpaeniformes | Platycephalidae | *Platycephalus indicus* (Linnaeus, 1758) | DD | Unknown |
| 153 | Gadiformes | Bregmacerotidae | *Bregmaceros mcclellandi* (Thompson, 1840) | NE | Unknown |
| 154 | Tetraodontiformes | Tetraodontidae | *Dichotomyctere fluviatilis* (Hamilton, 1822) | NE | Unknown |
